# The mirror neuron system compensates for amygdala dysfunction-associated social deficits in individuals with higher autistic traits

**DOI:** 10.1101/2021.05.17.444574

**Authors:** Lei Xu, Xiaoxiao Zheng, Shuxia Yao, Jialin Li, Meina Fu, Keshuang Li, Weihua Zhao, Hong Li, Benjamin Becker, Keith M. Kendrick

## Abstract

The amygdala is a core node in the social brain which exhibits structural and functional abnormalities in Autism spectrum disorder and there is evidence that the mirror neuron system (MNS) can functionally compensate for impaired emotion processing following amygdala lesions. In the current study, we employed an fMRI paradigm in 241 subjects investigating MNS and amygdala responses to observation, imagination and imitation of dynamic facial expressions and whether these differed in individuals with higher as opposed to lower autistic traits. Results indicated that individuals with higher compared to lower autistic traits showed reduced left amygdala responses to imitation and enhanced responses in the left superior temporal sulcus (STS) of the MNS to observation, imagination and imitation. Additionally, functional connectivity between the left amygdala and the left STS as well as some other MNS regions was increased in individuals with higher autistic traits, especially during imitation of fearful expressions. The amygdala-MNS connectivity significantly moderated autistic group differences on recognition memory for fearful faces and real-life social network indices, indicating that increased amygdala-MNS connectivity could diminish the social behavioral differences between higher and lower autistic trait groups. Overall, findings demonstrate decreased imitation-related amygdala activity in individuals with higher autistic traits in the context of increased cortical MNS activity and amygdala-MNS connectivity which may functionally compensate for amygdala dysfunction and social deficits. Training targeting the MNS may capitalize on this compensatory mechanism for therapeutic benefits in Autism spectrum disorder.

## 1 Introduction

Autism spectrum disorder (ASD) is characterized by impairments in social interaction, including impaired emotion processing (Uljarevic & Hamilton, 2013), imitation (Edwards, 2014) and empathy (Harmsen, 2019). As a core node of the social brain with established roles in emotion processing and many aspects of social cognition (Frith, 2007; Hennessey, Andari, & Rainnie, 2018), the amygdala exhibits both structural and functional abnormalities in ASD (Baron-Cohen et al., 2000; Hennessey et al., 2018). Amygdala volume is correlated with the severity of core ASD symptoms (Bellani, Calderoni, Muratori, & Brambilla, 2013; Daan van Rooij et al., 2018) and individuals with ASD display aberrant (hyperactive or hypoactive) amygdala activity in response to faces, especially fearful ones (Ashwin, Baron-Cohen, Wheelwright, O’Riordan, & Bullmore, 2007; Harms, Martin, & Wallace, 2010; Tottenham et al., 2013). Another key component of the social brain is the mirror neuron system (MNS) comprised primarily of the precentral gyrus, inferior parietal lobule (IPL), the inferior frontal gyrus (IFG) and the superior temporal sulcus (STS), which responds equivalently to observed and performed actions and may be of particular importance for key social behaviors such as language, empathy and imitation (Cattaneo & Rizzolatti, 2009; Iacoboni & Dapretto, 2006; Molenberghs, Cunnington, & Mattingley, 2012). While studies have reported decreased (Dapretto et al., 2006; Ramachandran & Oberman, 2006), or increased (Martineau, Andersson, Barthélémy, Cottier, & Destrieux, 2010; Wadsworth, Maximo, Donnelly, & Kana, 2018) or normal (Dinstein et al., 2010; Marsh & Hamilton, 2011) MNS activity in ASD, two meta-analyses have concluded that MNS hyperactivity is most consistently found (Chan & Han, 2020; Yang & Hofmann, 2016). It has been proposed that MNS dysfunction in ASD is mainly due to abnormal functional connectivity either within the MNS itself, such as in the IFG-STS automatic mimicry route (Hamilton, 2008), or between the MNS and other brain networks that regulate top-down control or social cognitive functions (Hamilton, 2013; Yates & Hobson, 2020).

Case studies in twins with equivalent focal bilateral basolateral amygdala-lesion due to Urbach-Weithe disease have reported that while one of them showed impaired facial fear recognition and social network size the other did not (Becker et al., 2012). Similar, differences between these twins have been reported for other important social behaviors which engage the amygdala such as emotional empathy (Hurlemann et al., 2010). More importantly, the twin who exhibited intact facial fear recognition and a normal-sized social network showed a corresponding enhanced activation in the MNS compared to her twin as well as healthy control subjects (Becker et al., 2012; Mihov et al., 2013). These findings suggest that the MNS may be able to functionally compensate for amygdala pathology and help alleviate impaired emotion processing and social interactions, which might indicate a potential therapeutic strategy to alleviate social deficits in autism. Indeed, previous studies have suggested that behavioral protocols such as Reciprocal Imitation Training (RIT) can improve social-communication skills in individuals with ASD (Ingersoll, 2010, 2012; Ingersoll & Schreibman, 2006) and increased activity in MNS regions, such as the IFG, are associated with improved social functioning in autism (Bastiaansen et al., 2011; Enticott et al., 2012). Given the lack of efficient treatments for social-emotional impairments in ASD, determining an MNS-based compensatory mechanism for amygdala deficits and associated social impairments may permit the design of novel interventions which specifically capitalize on this neurofunctional compensation.

Similar to other disorders, ASD is dimensional, with autistic personality traits varying continuously across the general population and clinical ASD residing at the upper extreme of this continuum (Constantino & Todd, 2003; Lundström et al., 2012). Autistic traits can indeed be measured in individuals with and without clinical ASD using the Autism Spectrum Quotient (ASQ - Murray, Booth, McKenzie, Kuenssberg, & O’Donnell, 2014). Healthy individuals with higher autistic traits have sub-clinical characteristics but remain able to cope with the social interaction demands of society in everyday life. In such healthy individuals white matter connectivity between the amygdala and the STS in the MNS is also associated with autistic traits (Iidaka, Miyakoshi, Harada, & Nakai, 2012). Thus, the MNS may compensate for amygdala dysfunction in healthy individuals with higher autistic traits, not only in clinical populations.

Against this background, the present study employed an fMRI paradigm using dynamic emotional faces to firstly establish the components of the MNS in a large sample of healthy young adults (n = 241). In order to provide the most accurate assessment of the components of the MNS in our subjects we not only identified regions responding equivalently to observation and imitation of face expressions, in line with most previous studies, but additionally we required equivalent responses when subjects simply imagined them. Next, we investigated MNS and amygdala responses to dynamic facial expressions in individuals with higher compared to lower autistic traits (n = 156). Autistic traits were assessed using the Chinese version of the ASQ which exhibits good internal consistency across healthy and clinical samples in Chinese populations (Zhang et al., 2016). In addition, social network indices and recognition memory for emotional faces were assessed. We hypothesized that individuals with higher autistic traits would exhibit reduced amygdala responses to fearful faces in particular and that there would be correspondingly enhanced MNS activity and strengthened functional connectivity between the amygdala and MNS. We further hypothesized that the magnitude of these potential compensatory changes in the MNS would be associated with better social functioning.

## 2 Methods

### 2.1 Participants

A total of 256 healthy Chinese adults (132 males) participated in the present study. All participants were free from current or past medical, neurological or psychiatric conditions and MRI contraindications, and were required to abstain from intake of caffeine, alcohol, nicotine, or other psychoactive substances during the 24 h prior to the experiment. The study was approved by the local ethics committee (Institutional Review Board, University of Electronic Science and Technology of China) and was in accordance with the latest revision of the Declaration of Helsinki. All participants signed written informed consent and received monetary compensation for their participation. Fifteen participants were excluded due to technical failures (incomplete data, n = 9), left-handedness (n = 4) or excessive head motion in more than one run (> 3 mm translation or 3° rotation; n = 2), leaving a total of 241 participants (123 males; 17-29 years old, mean ± SD age = 21.60 ± 2.30 years) for conjunction analysis. To investigate behavioral and brain differences between individuals with higher and lower autistic traits as well as potential compensatory mechanisms, participants from the highest and lowest 30% of the ASQ score distribution in the present sample were categorized as higher (ASQ scores >= 25; n=77, 42 males) and lower (ASQ scores<=18; n=79, 41 males) autistic groups respectively and subsequently used for between-group analyses (overviews of the exclusion criteria are provided in Figure S1).

### 2.2 Questionnaires

To control for potential confounding effects of mood alterations or clinical symptom loads often associated with ASD, such as depression and social anxiety, participants completed self-report questionnaires (Chinese versions) before MRI scanning, including the Positive and Negative Affect Scale – PANAS (Watson, Clark, & Tellegen, 1988), Liebowitz Social Anxiety Scale – LSAS (Liebowitz, 1987) and Beck Depression Inventory – BDI-II (Beck, Steer, & Brown, 1996). Individual levels of trait autism were measured by the Autism Spectrum Quotient – ASQ (Baron-Cohen, Wheelwright, Skinner, Martin, & Clubley, 2001). Participants additionally completed the Social Network Index – SNI (Cohen, Doyle, Skoner, Rabin, & Gwaltney, 1997) which contains three subscales, number of high-contact roles (reflecting social network diversity), number of people in social network (reflecting social network size) and number of embedded networks (reflecting social network complexity). To integrate the three subscales, composite SNI scores were calculated (Becker et al., 2012; see Supplementary method for the calculation).

### 2.3 Task paradigm

A total of 100 facial video clips (3s) from 20 actors (10 males) displaying happy, angry, fear, disgust, and cheek-blowing expressions were selected from an in-house database of 222 videos of 21 males and 16 females for high emotion expression recognition (>80%) as determined by pilot ratings from an independent sample of 25 subjects (14 males, age = 22.40 ± 0.42 years). Expression genuineness, emotion intensity, valence and arousal were also rated (9-point scale). The five expression sets were matched for genuineness. The cheek blowing expression set (which served as a control) was rated as medium intensity (4.25 ± 0.21), medium arousal (4.11 ± 0.10) and neutral valence (4.68 ± 0.07) and the four emotional face expression sets were matched for intensity, arousal, and valence distance from the neutral scale midpoint of 5 (see Table S1).

The MNS task (see Figure 1) was presented in an event related design via E-prime 2.0 (http://www.pstnet.com/eprime.cfm, Psychology Software Tools, Pittsburgh, Pennsylvania, USA). Participants firstly performed two passive observation runs, and each was comprised of 50 video clips. A white fixation cross on a black background was displayed between video clips for 2-5.6 s (0.6 s gap) as a jittered inter-trial interval and served as an implicit baseline for the analysis. Then two imagination and imitation runs were presented and each contained 50 trials. In each trial, a video clip was firstly displayed for 3 s and participants were instructed to imagine they were making the same facial expression as the one presented in the video they were viewing, followed by a red fixation cross for 3 s which indicated to participants that they should imitate the expression. Similarly, a white fixation cross was displayed for 2-5.6 s (0.6 s gap) before the red fixation cross as a jittered inter-stimuli interval, as well as after the red fixation cross as a jittered inter-trial interval, which served as an implicit baseline. A total of 100 video clips (20 per expression) were randomly allocated to two homogeneous runs and presented in the same pseudorandomized order for all participants. Each run commenced with a 20 s and ended with a 10 s blank screen to achieve a stable signal, resulting in a total duration of 370 s for each observation run and 710 s for each imagination and imitation run.

**Figure 1.**
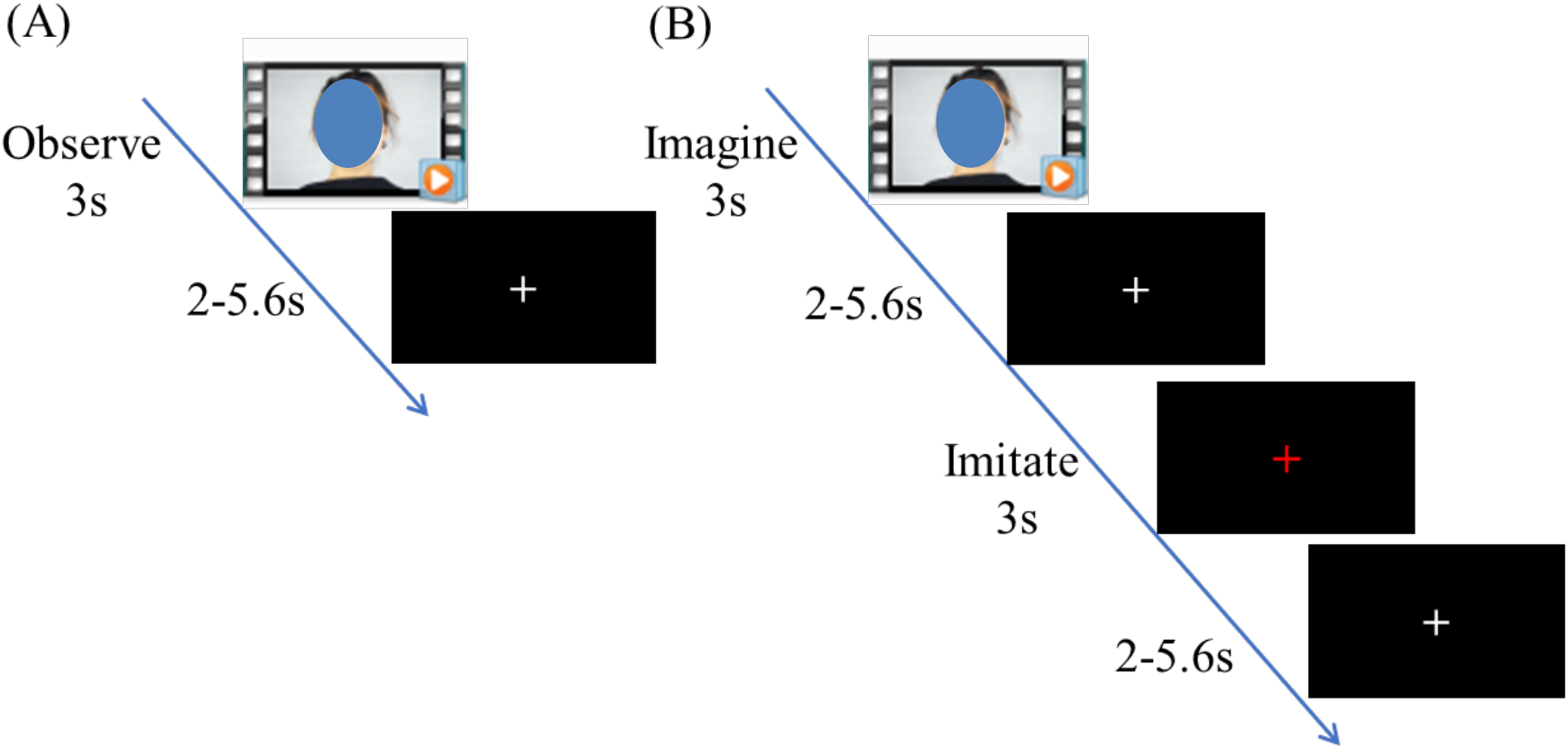
Example of a single trial in the MNS task. (A) Observation run; (B) imagination and imitation run. Note: for the preprint version that face has been overlayed with a blue oval.

After participants completed the MNS task, they were presented with static happy, angry, fearful, sad and neutral expressions (10 images for each expression with half male and half female) in the scanner during which they were only required to identify the gender of the individual. About forty min later, participants were given a memory recognition test where they were presented with the same face stimuli again together with an equal number of novel faces and asked to identify whether they were familiar or not (none of the faces were the same as in the MNS task).

### 2.4 Analytic approach for behavioral data

Questionnaire scores and recognition memory accuracy were not normally distributed (Shapiro-Wilk test, *ps*< 0.02; except for positive mood: *p* < 0.099) in the present sample and thus nonparametric analyses (Wilcoxon and Mann-Whitney U tests) were used to compare group differences between individuals with higher and lower autistic traits. Four subjects had incomplete data for Social Network Index and were excluded in analyses relating Social Network Index.

### 2.5 Analytic approach for neuroimaging data

#### 2.5.1 Image acquisition and data preprocessing

Neuroimaging data were acquired on a 3.0 Tesla GE Discovery MR750 system (General Electric Medical System, Milwaukee, WI, USA). Functional time-series were acquired using a T2*-weighted echo-planar imaging (EPI) sequence (repetition time: 2000 ms; echo time: 30 ms; flip angle: 90°; number of slices: 39 (interleaved ascending order); slice thickness: 3.4 mm; slice gap: 0.6 mm; field of view: 240 × 240 mm^2^; matrix size: 64 × 64). To improve normalization of the functional MRI data, high resolution T1-weighted structural images were additionally acquired using a 3D spoiled gradient recalled (SPGR) sequence (repetition time: 6 ms; echo time: minimum; flip angle: 9°; number of slices: 156; slice thickness: 1 mm without gap; field of view: 256 × 256 mm^2^; matrix size: 256 × 256). OptoActive MRI headphones (http://www.optoacoustics.com/) were used to reduce acoustic noise exposure for the participants during MRI data acquisition.

Preprocessing was conducted using standard procedures in SPM12 (Statistical Parametric Mapping, http://www.fil.ion.ucl.ac.uk/spm/) implemented in MATLAB 2014a (MathWorks, Inc., USA), including the following steps: (1) discarding the first 10 volumes of each run to allow MRI equilibration and active noise cancelling, (2) slice time correction, (3) head motion correction using a six-parameter rigid body algorithm, (4) tissue segmentation and skull-stripped biascorrection for the high-resolution structural images, (5) co-registration of mean functional images to structural images, (6) normalization (resampled at 3 × 3 × 3 mm) to Montreal Neurological Institute (MNI) space, and (7) spatial smoothing with an 8 mm full-width at half maximum (FWHM) Gaussian kernel. Runs with excessive head movement (>3 mm translation or >3° rotation) were excluded.

General linear model (GLM) analyses were implemented using SPM12. For each participant, fifteen condition-specific regressors (5 expressions x 3 task demands) were modeled using a stick function convolved with the canonical hemodynamic response function (HRF). The six head motion parameters were additionally included as nuisance regressors and a 128 s high-pass filter was applied to further control low-frequency noise artifacts.

#### 2.5.2 Whole brain analyses and thresholding

To determine MNS activation maps, full factorial F test and conjunction analysis based on the entire sample were conducted using a whole-brain approach with a conservative significance threshold of *p* < 0.05 peak level Family-Wise Error (FWE) correction and a minimum voxel size of *k* > 10. Brain regions were identified using the Automated Anatomic Labelling (AAL) atlas (Tzourio-Mazoyer et al., 2002) as implemented in the WFU Pick Atlas ((Maldjian, Laurienti, Kraft, & Burdette, 2003).

To investigate autism trait-related neurofunctional alterations and potential compensatory mechanisms in amygdala and MNS neural activity, individuals with higher and lower autistic traits were compared. Flexible factorial ANOVA models were created to examine three-way interaction effects involving the factors group (higher versus lower autism) x expression x task demand, as well as two-way interaction effects involving the factors group x expression or group x task demand, respectively. To robustly determine main effects of autistic trait group and to further disentangle significant interaction effects via post hoc comparisons, 2 sample t tests including sex, age, depression and social anxiety as covariates were employed. These analyses employed a whole-brain approach with a significance threshold of *p* < 0.05 cluster level FWE corrected for multiple comparisons. In line with recommendations for the control of false-positives rates in cluster based correction methods an initial cluster defining threshold of *p* < 0.001 uncorrected was applied (Eklund, Nichols, & Knutsson, 2016; Slotnick, 2017). Given the differential functional roles of amygdala subregions in social-emotional processes amygdala effects were mapped to probabilistic amygdala subregions using the Anatomy toolbox (Amunts et al., 2005; Eickhoff et al., 2005).

A generalized psychophysiological interactions (gPPI) (McLaren, Ries, Xu, & Johnson, 2012) analysis was implemented to investigate network level differences between the higher and lower autistic trait groups on the level of task-modulated functional connectivity. To this end, seed regions were determined on the basis of significant results from the BOLD level group difference analyses using the identified clusters as seed regions. Group differences in seed-to-whole brain functional connectivity were examined using 2 sample t test with the contrasts showing significant results from the BOLD level group difference analyses. Again, sex, age, depression and social anxiety were included as covariates. Based on our a priori hypothesis focusing on the interaction between the amygdala and MNS the gPPI analyses focused on the MNS network employing a significance threshold of *p* < 0.05 peak level FWE adopted for the MNS using a small volume correction (SVC) approach. The corresponding MNS mask was created by combining anatomically defined core regions of the MNS (STG, IFG and IPL, from the AAL atlas) with the study specific MNS activity pattern (conjunction activity map for the emotional MNS).

#### 2.5.3 Confirmatory region of interest (ROI) analyses

Region of interest (ROI) analyses were further performed to confirm whether the MNS regions exhibiting compensatory changes following amygdala damage in previous studies also showed enhanced responses in individuals with higher autistic traits. ROIs were selected based on previous studies (Becker et al., 2012; Mihov et al., 2013) as 6mm radius spheres centered at the following coordinates (MNI space): left IFG (−48, 6, 28) and left STS (−58, −44, 22), left premotor cortex face area (−32, 6, 63; corresponding Talairach space at −32, 9, 58), as well as the anatomically defined bilateral IPL (AAL atlas).

For activation, parameter estimates of signal intensity in these brain regions were extracted by MarsBaR (Brett, Anton, Valabregue, & Poline, 2002) and then subjected to repeated measures ANOVAs with task demand and expression as within-subjects factors and autistic trait group as a between-subjects factor.

For connectivity, parameter estimates of connectivity strength from gPPI analysis in predefined ROIs which exhibited compensatory changes (increased activity) in the higher autistic trait group were further extracted and subjected to repeated measures ANOVAs.

#### 2.6 Analytic approach for associations between brain and behavioral alterations

To examine associations between brain and behavioral alterations, individual parameter estimates of signal intensity or functional connectivity in the brain regions which showed significant group differences were extracted. Exploratory moderation models were implemented to examine associations between autistic trait groups, brain and behavioral alterations, specifically whether the brain activity or connectivity moderated the autistic trait group differences on behavioral indices. Separate models were tested for each moderator (activity or connectivity), as well as each behavioral outcome. We estimated the moderation effects using the PROCESS macro for SPSS (Model 1) (Hayes, 2013). The Johnson-Neyman method was additionally used to calculate the entire range of moderator variable values (ie. amygdala activation) for which the focal predictor (i.e., autistic groups) is significantly associated with the dependent variable (i.e., social network size).

## 3 Results

### 3.1 Behavioral results

Wilcoxon and Mann-Whitney U tests revealed that individuals with higher autistic traits displayed significantly decreased positive mood and increased negative mood, as well as increased symptom loads of depression and social anxiety compared with the lower trait group (see Table 1). Nonetheless, individuals with higher autistic traits remained within minimal depression (0-13; Beck et al., 1996) and mild to moderate social anxiety (<55 mild, 55-65 moderate; Liebowitz, 1987). Notably, recognition memory for fearful faces and social network indices were significantly decreased in individuals with higher autistic traits (all *ps* < 0.047). Full details on the recognition memory accuracy and social network indices are provided in Table 1.

**Table 1.** Age, questionnaire scores and behavioral performance for higher and lower autistic trait groups (mean ± sem)

| Behavior measures | Overall<br>(n=241) | Higher group<br>(n=77) | Lower group<br>(n=79) | *p* |
| --- | --- | --- | --- | --- |
| Age | 21.60±0.15 | 21.75±0.28 | 21.66±0.26 | 0.997 |
| Autism Spectrum Quotient | 21.42±0.37 | 27.99±0.31 | 15.01±0.28 | <0.001 |
| Beck Depression Inventory | 8.37±0.48 | 10.55±0.92 | 5.89±0.69 | <0.001 |
| Liebowitz Social Anxiety Scale | 44.62±1.54 | 55.42±2.81 | 33.70±2.25 | <0.001 |
| Positive and Negative Affect Scale |  |  |  |  |
| -positive | 28.63±0.43 | 26.32±0.70 | 31.57±0.68 | <0.001 |
| -negative | 17.62±0.39 | 18.38±0.72 | 16.52±0.69 | 0.049 |
| Memory accuracy (%) |  |  |  |  |
| -happy | 58.73±0.76 | 57.21±1.40 | 59.05±1.38 | 0.266 |
| -angry | 55.17±0.68 | 54.81±1.26 | 55.32±1.25 | 0.698 |
| -fearful | 62.53±0.68 | 60.65±1.41 | 64.18±0.98 | 0.047 |
| -sad | 60.23±0.63 | 60.91±1.14 | 60.70±1.05 | 0.764 |
| -neutral | 55.89±0.64 | 57.66±1.26 | 55.89±0.99 | 0.382 |
| Social Network Index | n=237 | n=76 | n=77 |  |
| -N of high contact roles | 3.62±0.94 | 3.38±0.19 | 3.88±0.17 | 0.039 |
| -N of high contact people | 11.75±0.55 | 9.91±0.95 | 13.65±0.99 | 0.001 |
| -N of embedded networks | 0.93±0.07 | 0.71±0.11 | 1.12±0.12 | 0.003 |
| Overall composite score | 99.94±0.65 | 97.68±1.18 | 102.15±1.12 | 0.001 |

Notes: Four subjects had incomplete data for Social Network Index and were excluded in analyses relating Social Network Index.

### 3.2 Neuroimaging results

#### 3.2.1 MNS activation

In an initial step the MNS was mapped by identifying regions commonly activated during observation, imagination and imitation of the facial expressions. To this end conjunction analyses using the conjunction-null hypothesis for the specific contrasts, including observation > baseline, imagination > baseline and imitation > baseline across all conditions (general MNS) or across the four emotional expressions (emotional MNS) were calculated respectively. Commonly activated regions across all facial expressions included typical MNS regions including the IPL, STS, IFG and precentral gyrus, but also middle temporal gyrus (MTG), fusiform gyrus and cerebellum, supplementary motor area (SMA) and postcentral gyrus. The emotional MNS (only responsive to emotional faces) showed a comparable pattern than the general one but notably extended to the bilateral amygdala, insula and thalamus (see Figure 2 and Table S2 for whole-brain analyses with a significance threshold of *p* < 0.05 peak level FWE corrected, *k* > 10).

**Figure 2.**
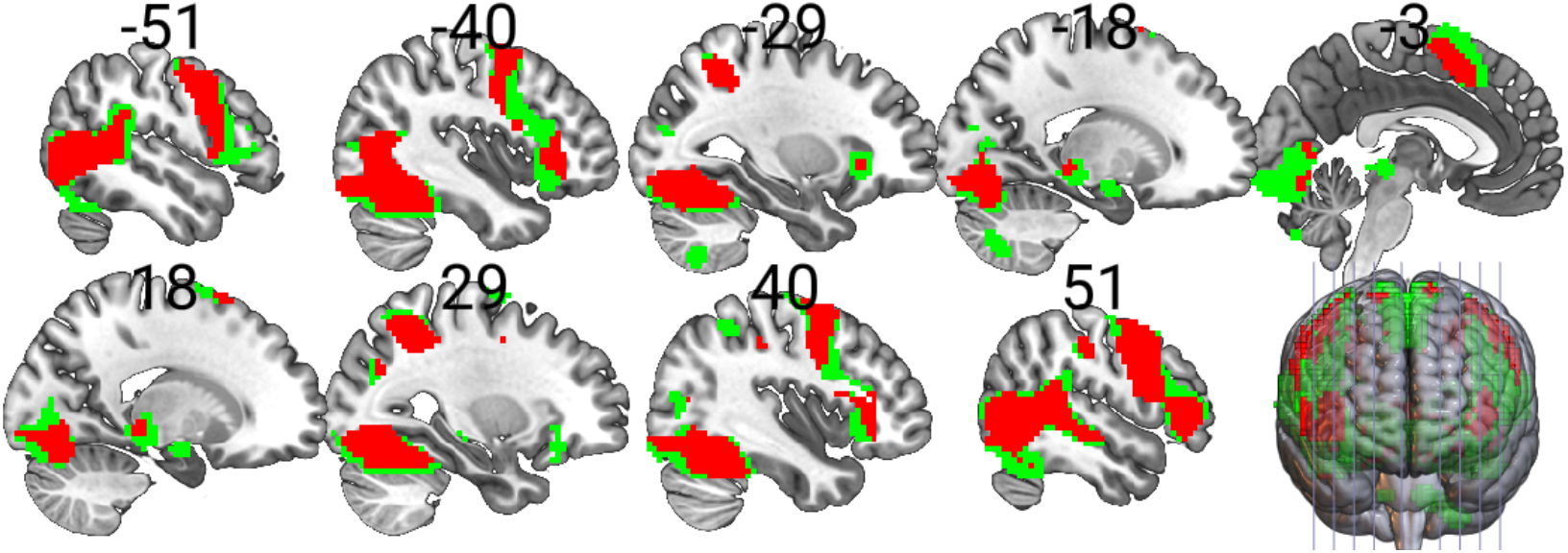
Conjunction map of observation, imagination and imitation across the five facial expressions (red, conjunction across 15 conditions, *p* < 0.05 peak level FWE corrected, *k* > 10) and across the four emotional facial expressions (red and green, conjunction across 12 emotional conditions, *p* < 0.05 peak level FWE corrected, *k* > 10).

#### 3.2.2 Differences in neural responses between lower and higher trait autism groups

The overall main effect of autistic trait group on brain activity examined by 2 sample t-test with sex, age, depression and social anxiety as covariates revealed no significant group differences arguing against unspecific effects of trait autism on brain activation. No significant interaction effects of group x expression x task demand and group x expression on brain activity were observed in the flexible factorial ANOVA models. However, a significant group x task demand interaction effect on brain activity was found in the left amygdala (whole brain analysis: *x, y, z* = −27, −10, −31; *F*_2,308_ = 12.31, *k* = 109; *p* = 0.021 cluster level FWE corrected; see figure 3A). The significant interaction effect was further examined by task demand-specific post hoc voxel-wise two sample t-tests comparing the groups for observation, imagination and imitation independently while controlling for sex, age, levels of depression and social anxiety as covariates. Between group differences were specifically observed during imitation, such that individuals with higher autistic traits exhibited significantly decreased activation as compared to those in the lower autistic trait group in the left amygdala (whole brain analysis: *x, y, z* = −24, −7, −22; *T*_150_ = 4.68, *k* = 276; *p* = 0.001 cluster level FWE corrected; see figure 3B). According to the Anatomy toolbox (Amunts et al., 2005; Eickhoff et al., 2005), the cluster primarily encompassed the basolateral subregion of the amygdala (BLA, 73.3% of the probabilistic BLA mask overlapped with the cluster and the peak voxel located with 80% probability in the BLA). No significant group difference was found during observation or imagination.

**Figure 3.**
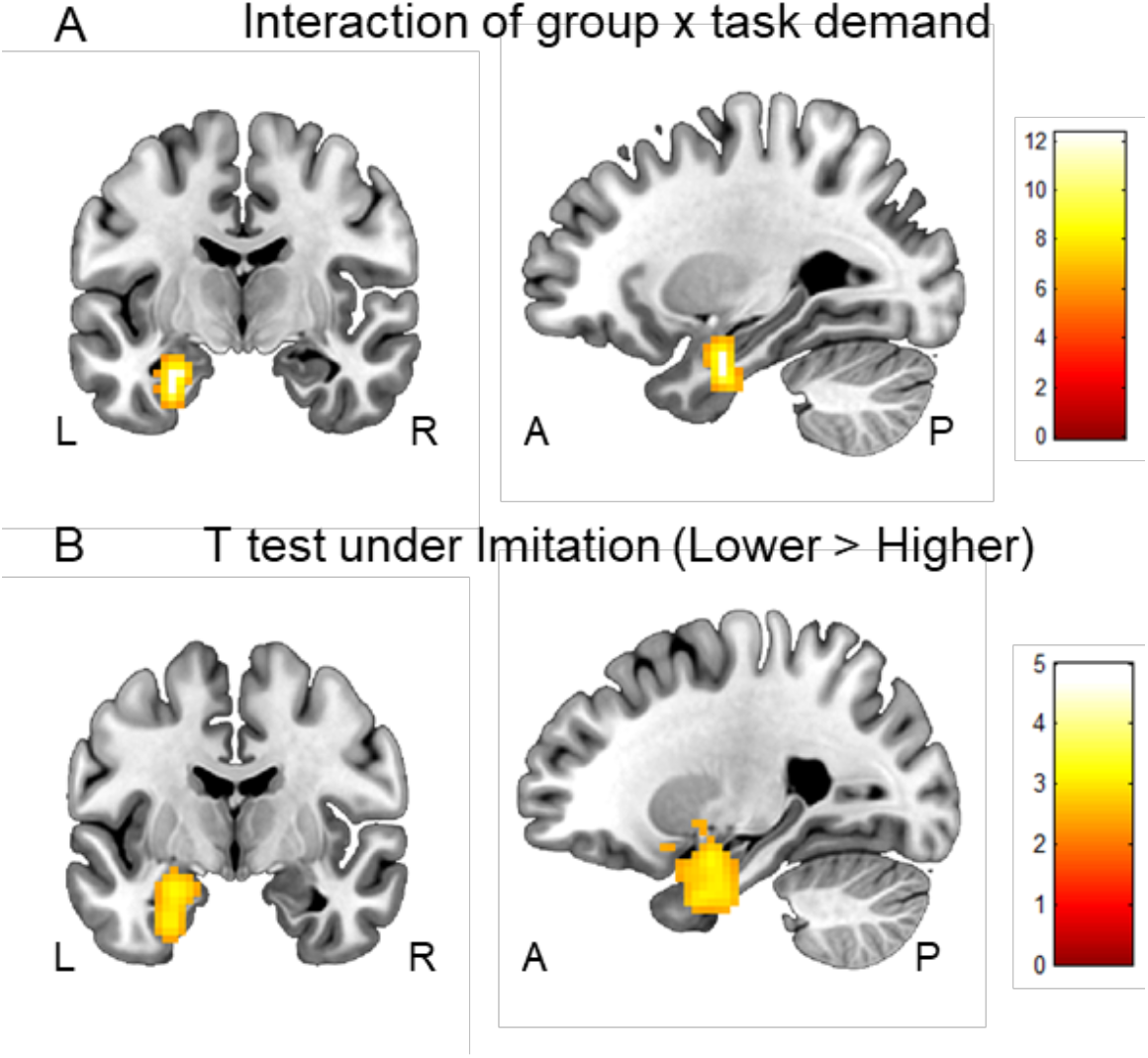
Group differences in brain activity. (A) Interaction of group x task demand in the left amygdala (*x, y, z* = −27, −10, −31; *p* = 0.021 cluster level FWE correction; displayed at threshold: _*p*uncorr_ < 0.001). (B) The higher autistic trait group showed significantly decreased activity in the left amygdala only during imitation compared with the lower autistic trait group (−24, −7, −22; *p* = 0.001 cluster level FWE correction; displayed at threshold: _*p*uncorr_ < 0.001).

#### 3.2.3. Differences in functional connectivity between lower and higher trait autism groups

Based on brain activity findings, gPPI analyses were further conducted using the cluster in the left amygdala that exhibited significant BOLD activity differences between the groups as a seed region. Given that group differences in neural activity were specifically observed during imitation, the examination of group differences on functional connectivity focused on the imitation condition and examined each emotional expression separately, using two sample t tests with sex, age, levels of depression and social anxiety being controlled for. Whole-brain analyses suggested that when imitating fearful expressions, the higher autistic trait group exhibited relatively increased functional connectivity between the left amygdala and core MNS regions, specifically the right STS (*x, y, z* = 57, −31, 11; *T*_150_ = 3.77, *k* = 34, *p* = 0.041 SVC peak level FWE corrected) and the left IFG (*x, y, z* = −42, 26, 2; *T*_150_ = 3.72, *k* = 38; *p* = 0.048 SVC peak level FWE corrected; see figure 4A). Results remained stable if the seed region of the activated cluster overlapped with the amygdala template (created using probabilistic maps from the Anatomy toolbox), suggesting a robust contribution of the amygdala(see Supplementary results and Figure S2). No significant group differences were found under imitation of other expressions.

**Figure 4.**
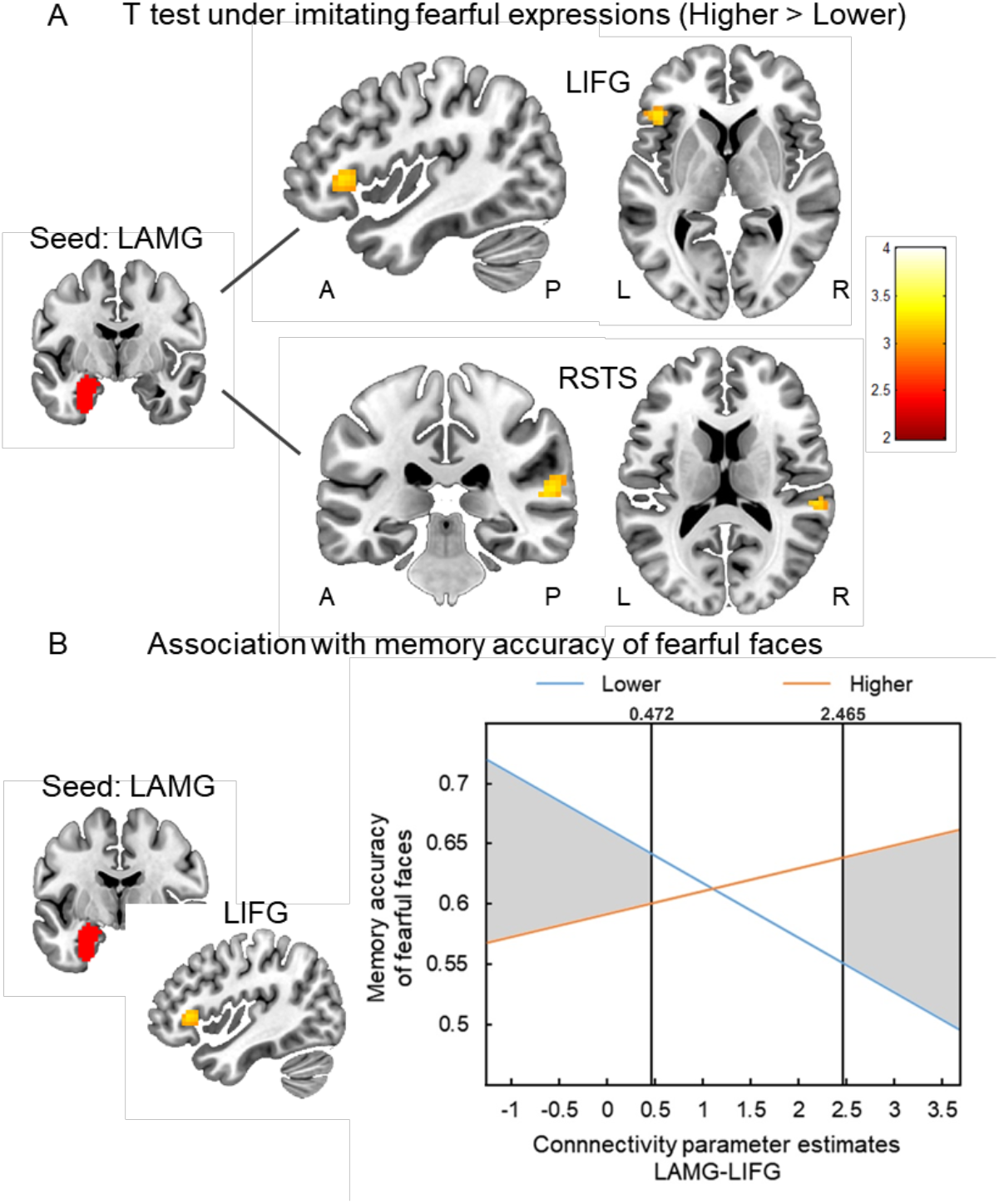
Group differences in functional connectivity between the left amygdala and the MNS, as well as the associations with behavioral memory accuracy of fearful faces. (A) Individuals in the higher autistic trait group exhibited increased functional connectivity of the left amygdala (LAMG: −24, −7, −22; seed region) to the right superior temporal sulcus (RSTS: 57, −31, 11) and left inferior frontal gyrus (LIFG: −42, 26, 2) during imitating fearful expressions (*p* < 0.05 SVC peak level FWE correction). (B) LAMG-LIFG connectivity during imitation of fearful faces moderated the autistic group differences on memory accuracy of fearful faces. Shaded area identifies regions of significance in which memory accuracy of the fearful face differs significantly between the higher and lower autistic trait groups at *p* < 0.05. The outer border represents the lowest and highest connectivity parameter estimates in the sample.

A moderation analysis revealed a significant moderation effect (R^2^ = 0.3024, *F*_7,148_ = 2.1277, *p* = 0.0441) after controlling for sex, age, levels of depression and social anxiety, reflecting that connectivity of left amygdala and left IFG under imitating fearful faces significantly moderated the association between trait autism groups and memory accuracy of fearful faces (B = 0.0644, S.E. = 0.0228, *T*_148_ = 2.8284, *p* = 0.0053). Further disentangling the effect using the Johnson-Neyman technique revealed that autistic trait group differences on memory accuracy of fearful faces would be significant when the connectivity was below 0.4722 (40.38%; lower > higher) or above 2.4651 (96.15%; higher >lower; see figure 4B). Thus, increased left amygdala-IFG coupling during imitation of fearful faces could diminish and even reverse the behavioral differences on memory accuracy of fearful faces between higher and lower autistic trait groups. Moderation analysis for connectivity of left amygdala and right STS under imitating fearful faces as well as activation of left amygdala under imitation failed to reach significance (see Table S3).

#### 3.2.4 ROI-based analysis of MNS regions which may compensate for amygdala deficits

Examination of the regions which have been reported as potential compensatory regions facilitating MNS-mediated compensation of amygdala deficits (Becker et al., 2012; Mihov et al., 2013) were tested by means of extracted activity-based parameter estimates from predefined ROIs. Repeated measures ANOVAs revealed a significant main effect of autistic trait group in the left STS (−58, −44, 22) such that the higher autistic trait group exhibited increased activity across all conditions (i.e. observation, imagination and imitation) (*F*_1,154_ = 4.341, *p* = 0.039, *η*^2^_p_ = 0.027; see figure 5A). No significant main effects of autistic trait group or interactions with autistic trait group were found for other ROIs (all *ps* > 0.160).

**Figure 5.**
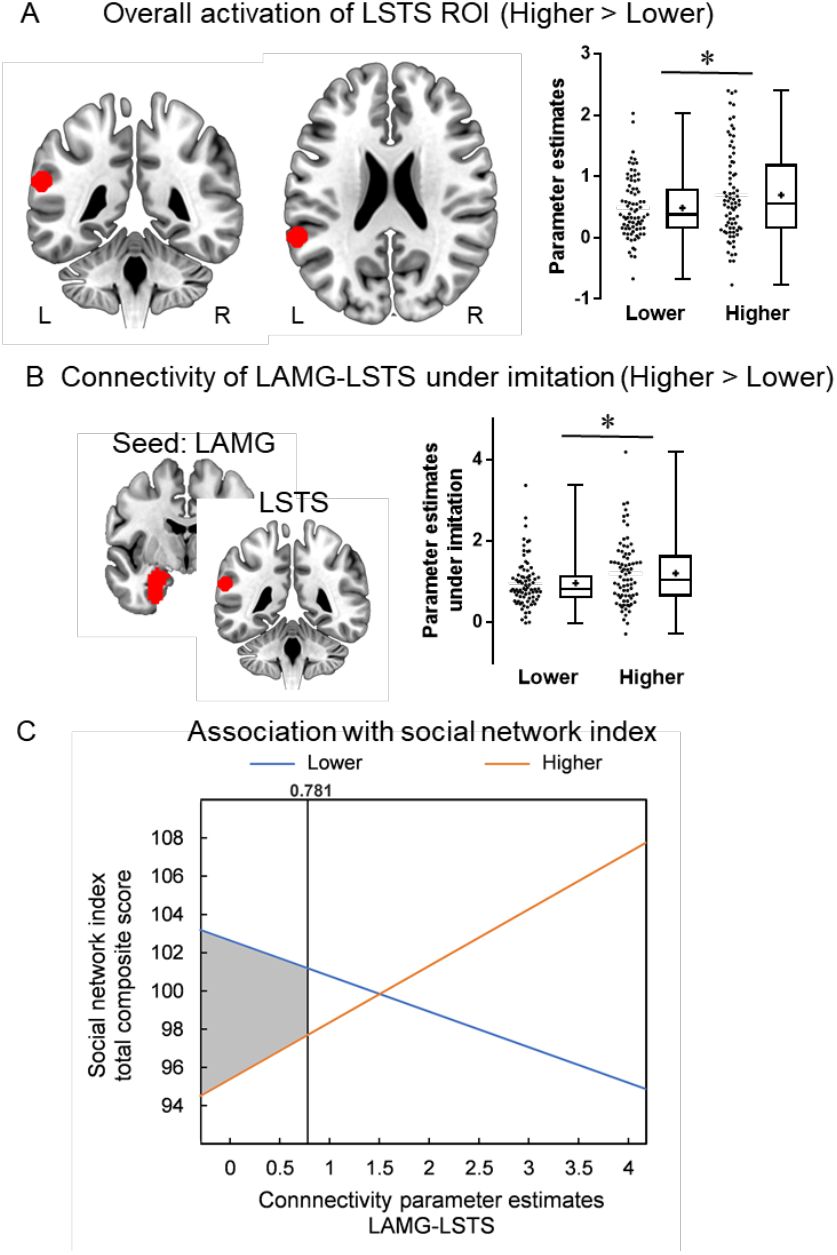
Results of ROI-based analysis. (A) The higher autistic trait group showed significant increased activity in the ROI of left superior temporal gyrus (LSTS: −58, −44, 22) across all conditions (i.e. observation, imagination and imitation) compared with the lower autistic trait group (\**p* < 0.05). (B) The higher autistic trait group showed significant increased left amygdala (LAMG: −24, −7, −22; seed region) connectivity with the left STS ROI (−58, −44, 22) during imitation compared with the lower trait autism group (\**p* < 0.05). (C) LAMG-LSTS connectivity during imitation moderated the autistic group differences on social network size. Shaded area identifies regions of significance in which social network size differs significantly between the higher and lower autistic trait groups at *p* < 0.05. The outer border represents the lowest and highest connectivity parameter estimates in the sample.

Next, extracted connectivity-based parameter estimates (left amygdala as seed region) of the predefined left STS ROI (−58, −44, 22) under the imitation condition were subjected to repeated measures ANOVA with expression as within-subjects factor and autistic trait group as a between-subjects factor. A significant main effect of autistic trait group indicated that the higher autistic trait group exhibited increased functional connectivity between left amygdala and left STS while imitating facial expressions (*F*_1,154_ = 4.596, *p* = 0.034, *η*^2^_p_ = 0.029; see figure 5B).

A moderation analysis revealed a significant moderation effect (R^2^ = 0.2270, *F*_7,145_ = 6.0822, *p* < 0.001) after controlling for sex, age, levels of depression and social anxiety, reflecting that left amygdala connectivity with the left STS under overall imitation condition significantly moderated the association between trait autism groups and composite SNI score (B = 4.8193, S.E. = 2.3256, *T*_145_ = 2.0723, *p* = 0.0400). Further disentangling the effect using the Johnson-Neyman technique revealed that autistic group differences on memory accuracy of fearful faces would be significant when the connectivity was below 0.7807 (38.56%; lower > higher; see figure 5C). Thus, increased connectivity of left amygdala and left STS during imitation could diminish the social network size differences between higher and lower autistic trait groups. Moderation analysis for activation of the predefined left STS ROI across all conditions failed to reach significance (see Table S4).

## 4 Discussion

The present fMRI study aimed firstly to comprehensively establish the MNS in a large cohort of healthy adults using a conjunction analysis of regions responding equivalently to observation, imagination and imitation of classical dynamic emotional and non-emotional control face expressions and then investigate whether and how the MNS may compensate for amygdala and social dysfunction in individuals with higher as opposed to lower autistic traits. Results of the conjunction analysis in the entire sample revealed that core components of the MNS, including the IFG, precentral gyrus, IPL and STS all responded equivalently to both emotional and control expressions whereas subcortical regions such as the amygdala and thalamus only did so for classical emotional expressions. Individuals in the sub-group with higher trait autism scores exhibited worse recognition memory for fearful faces and smaller real-life social networks at the behavioral level, whereas at the neural level they showed decreased left amygdala activity during imitation and increased left STS activity during observation, imagination and imitation. Furthermore, functional connectivity between these two regions was stronger in the higher autistic trait group during imitation and moderated the autistic group differences on social network scores. Left amygdala connectivity with right STS and left IFG during imitation of fearful expressions were also stronger in higher autistic group and amygdala-IFG connectivity moderated the autistic group differences on recognition memory accuracy of fearful faces. Overall, our findings indicate that increased amygdala-MNS connectivity could diminish social behavior differences between higher and lower autistic trait groups.

In terms of establishing the extent of the MNS in the human brain the conjunction analysis carried out in the current study including imagination as well as observation and imitation is the largest single one to date. Importantly it largely replicates previous findings reported in a metaanalysis of 125 studies (Molenberghs et al., 2012) in confirming the inclusion of not only the most commonly reported regions (IFG, STS, IPL and precentral gyrus) but also others such as the cerebellum, fusiform gyrus, hippocampus, medial temporal gyrus, supplementary motor area and superior occipital lobe. Additionally, sub-cortical regions such as the amygdala, insula and thalamus exhibited mirror properties, but only for emotional expressions and form part of what could be termed an emotional MNS (van der Gaag, Minderaa, & Keysers, 2007) and has been suggested to integrate motor mimicry with emotional aspects to facilitate social-emotional functions such as facial emotion recognition or empathy.

Our findings that healthy individuals with higher autistic traits exhibited worse recognition memory for fearful faces and smaller real-life social networks compared to those with lower autistic traits are consistent with previous findings in ASD patients (Lozier, Vanmeter, & Marsh, 2014; Uljarevic & Hamilton, 2013; van Asselt-Goverts, Embregts, Hendriks, Wegman, & Teunisse, 2015) and further confirmed the validity of dimensional approach to ASD. In line with the amygdala theory of autism (Baron-Cohen et al., 2000), our findings indicate that healthy individuals with higher autistic traits display hypoactivation in the left amygdala during imitation of facial expressions. Most studies in ASD patients have also reported reduced amygdala activation during dynamic or static face processing (Ashwin et al., 2007; Bölte et al., 2018; Ciaramidaro et al., 2018; Hadjikhani, Joseph, Snyder, & Tager-Flusberg, 2007; Harms et al., 2010; Pelphrey, Morris, McCarthy, & LaBar, 2007), as well as a previous Activation Likelihood Estimation (ALE) meta-analysis in ASD (Di Martino et al., 2009). However, some other studies have reported amygdala hyperactivity in ASD, although this was more related to comorbid anxiety symptoms (Herrington, Miller, Pandey, & Schultz, 2016; Kleinhans et al., 2010) or attention to eyes (Dalton et al., 2005; Hadjikhani et al., 2017; Lassalle et al., 2017; Monk et al., 2010; Tottenham et al., 2013). Thus, it was not surprising that the current study showed amygdala hypoactivity in a free viewing paradigm while controlling for social anxiety. The present findings coupled with previous findings in ASD proved the amygdala dysfunction as a neurobiological marker of autism and confirm the potential therapeutic relevance of targeting neurofunctional mechanism that can compensate for amygdala dysfunction and associated social deficits in ASD.

Similar to an amygdala-damaged patient with intact fear recognition and social network size (Becker et al., 2012; Mihov et al., 2013), healthy individuals with higher autistic traits showed MNS hyperactivity in the same location of the left STS. Although consistent with previous findings in ASD patients (Chan & Han, 2020; Yang & Hofmann, 2016), the current finding of increased STS activation in individuals with subclinical autistic characteristics but intact social interaction indicates the possibility of functional MNS compensation for amygdala pathology in autistic individuals.

More importantly, our present study further found increased functional connectivity of left amygdala and MNS regions, including bilateral STG and left IFG, when individuals with higher autistic traits imitated (fearful) face expressions, and these connectivity increases could diminish social behavior differences between higher and lower autistic trait groups, such as recognition memory for fearful faces and personal social network size. Previous studies in ASD patients have reported weaker functional connectivity between the amygdala and superior/medial temporal lobe or inferior/medial frontal lobes, and the connectivity was significantly correlated with autistic symptom severity (Abrams et al., 2013; Monk et al., 2010; Shen et al., 2016). However, in individuals with higher autistic traits, we found increased connectivity of the left amygdala and MNS regions. This is consistent with a structural study in healthy young adults which showed a positive correlation between autistic traits and white matter connectivity of the amygdala and STS (Iidaka et al., 2012). Furthermore, the present finding that the increased amygdala-MNS connectivity was associated with better recognition memory for fearful faces and social network size further indicates the possibility of MNS compensation for social deficits in healthy individuals with higher autistic traits. Interestingly, individuals with ASD are slower and less precise at imitating facial expressions and the better they are imitating them the more accurate they are at recognizing face expressions (Drimalla, Baskow, Behnia, Roepke, & Dziobek, 2021). This together with our findings showing that the influence of higher autistic traits on amygdala functional connectivity with the MNS is particularly in the context of imitation would seem to suggest that imitation training might be an effective therapeutic strategy. Indeed, several studies have reported positive effects of imitation training in ASD (Ingersoll, 2010, 2012; Ingersoll & Schreibman, 2006).

The current study has several limitations. Firstly, only face-related stimuli were used for observation, imagination and imitation and not of other parts of the body such as the hands. Secondly, we did not make any assessments of the quality of the imitation exhibited by subjects which might have been associated with observed behavioral and neural differences in the lower and higher trait autism groups.

Taken together, the present study demonstrates that the MNS may be able to compensate for amygdala dysfunction and social deficits in healthy individuals with higher autistic traits. Thus, MNS based training may be of potential therapeutic benefit for ASD. Since ASD has different subtypes, it is possible that effective compensation in the MNS may occur more in some subtypes, such as high functioning autism, Asperger’s syndrome and broad autism phenotypes. Subtyping has also been proposed on the basis of different patterns of social behavior (Aloof, Passive and Active-but-odd – Meng et al., 2018), which might also potentially reflect differential levels of amygdala-MNS functional compensation. Further work is therefore needed to explore functional compensation in the MNS in different subtypes of ASD, including in children and adolescents, and to observe potential beneficial effects of MNS training regimes such as face expression imitation.

## Supplementary Information

### Methods

In order to integrate the three subscales of the Social Network Index (SNI) in a single measure, composite SNI scores were calculated in three steps: first, individual scores in each subscale were z-standardized within the entire sample; second, z-scores from the three subscales were added for each individual; third, sum scores were z-standardized and transformed to a normal distribution with a mean of 100 and a standard deviation of 10, resulting in the final composite scores.

### Results

The cluster that exhibited significant group differences on activation were overlapped with amygdala template which combined probabilistic maps (thresholded at > 50% probability) of basolateral, centromedial, and superficial amygdala subregions from the Anatomy toolbox and then served as the seed region for gPPI analysis. Whole-brain results of two sample t test after controlling sex, age, levels of depression and social anxiety under the condition of imitating fearful expressions revealed significant and stable group differences on the amygdala connectivity with the right STS (*x, y, z* = 63, −40, 20; *T_150_* = 3.66, *k* = 23, *p* = 0.056 SVC peak level FWE corrected) and the left IFG (*x, y, z* = −42, 26, 2; *T_150_* = 3.79, *k* = 33; *p* = 0.037 SVC peak level FWE corrected; see figure S2).

## Supplementary Figures

**Figure S1.**
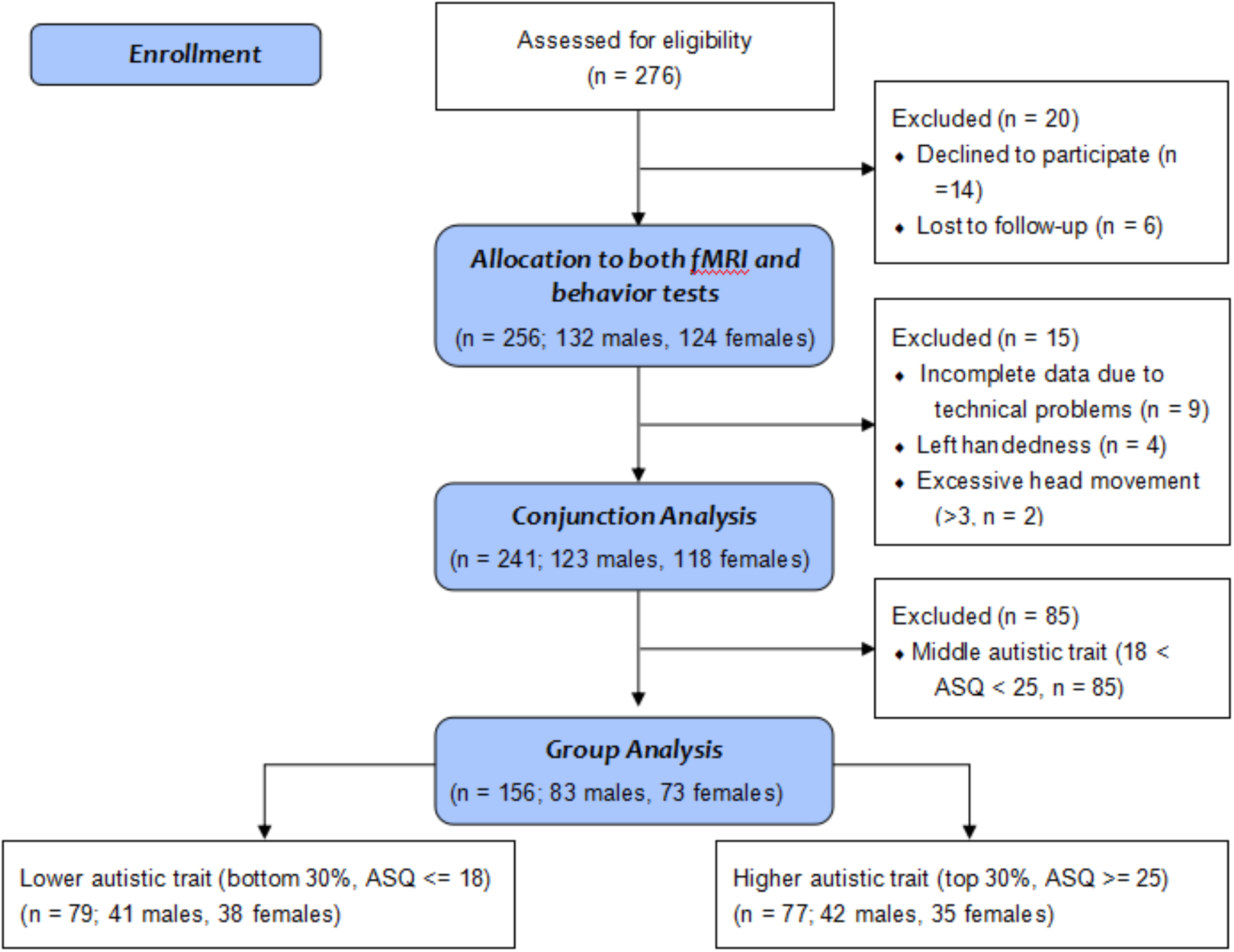
Flow diagram displaying exclusion of participant and rationale for exclusion.

**Figure S2.**
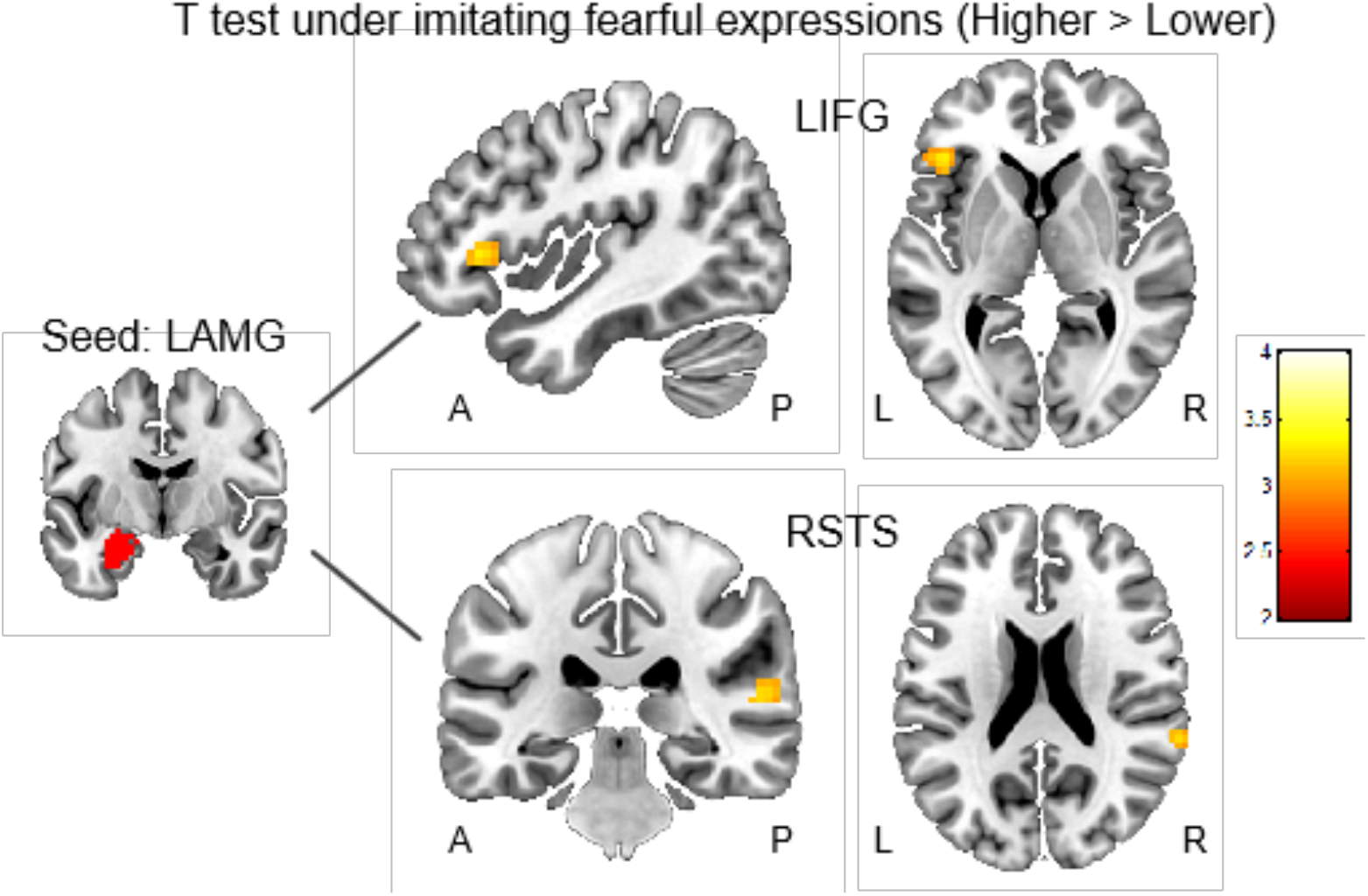
Whole brain results of group differences in functional connectivity of left amygdala (LAMG: the overlapped region of the activated cluster (−24, −7, −22) and amygdala template; seed region). Individuals in the higher autistic trait group exhibited increased left amygdala connectivity with the right superior temporal sulcus (RSTS: 63, −40, 20; *p* = 0.056 SVC peak level FWE corrected) and left inferior frontal gyrus (LIFG: −42, 26, 2; *p* = 0.037 SVC peak level FWE corrected) during imitating fearful expressions.

## Supplementary Tables

**Table S1.**
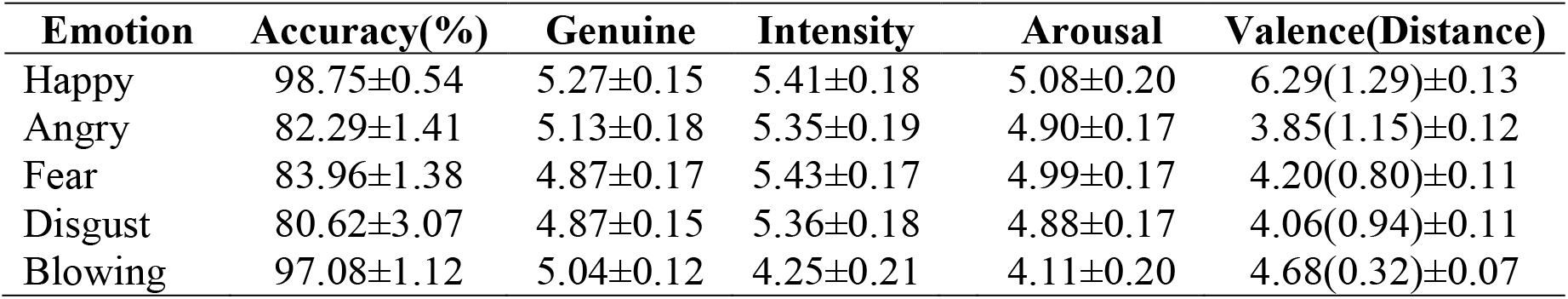
Pilot rating for video clips of each expression (mean ± sem)

**Table S2.** Common activated areas under observation, imagination and imitation.

| Region | Side | $k$ | MNI coordinates | | | $t$ | $P_{\text{FWE}}$ |
| --- | --- | --- | --- | --- | --- | --- | --- |
| | | | $x$ | $y$ | $z$ | | |
| Conjunction across the five facial expressions |  |  |  |  |  |  |  |
| Precentral Gyrus | R | 888 | 48 | 5 | 35 | 22.62 | < 0.001 |
| Precentral Gyrus | R |  | 54 | 8 | 41 | 22.50 |  |
| Triangular Inferior Frontal Gyrus | R |  | 51 | 32 | 2 | 14.23 |  |
| Fusiform/Cerebellum 6 | R | 3335 | 30 | -61 | -19 | 22.60 | < 0.001 |
| Fusiform/Cerebellum 6 | L |  | -33 | -55 | -22 | 22.14 |  |
| Middle Temporal Gyrus | L |  | -48 | -61 | 5 | 21.93 |  |
| Precentral Gyrus | L | 479 | -51 | 5 | 47 | 18.10 | < 0.001 |
| Precentral Gyrus | L |  | -42 | -1 | 32 | 13.30 |  |
| Rolandic Operculum | L |  | -48 | 8 | 2 | 5.72 |  |
| Inferior Parietal Lobule | R | 181 | 30 | -49 | 50 | 14.98 | < 0.001 |
| Superior Occipital Lobule | R |  | 27 | -70 | 32 | 6.57 |  |
| Supplementary Motor Area | R | 341 | 6 | 11 | 53 | 14.15 | < 0.001 |
| Supplementary Motor Area | R |  | 15 | 17 | 68 | 6.08 |  |
| Inferior Parietal Lobule | L | 99 | -27 | -52 | 47 | 8.67 | < 0.001 |
| Hippocampus | R | 22 | 21 | -31 | -7 | 8.61 | < 0.001 |
| Triangular Inferior Frontal Gyrus | L | 109 | -39 | 29 | -1 | 7.62 | < 0.001 |
| Hippocampus | L | 13 | -18 | -31 | -4 | 7.35 | < 0.001 |
| Postcentral Gyrus | R | 77 | 48 | -25 | 41 | 6.69 | < 0.001 |
| Postcentral Gyrus | R |  | 57 | -16 | 35 | 5.96 |  |
| Conjunction across the four emotional facial expressions |  |  |  |  |  |  |  |
| Precentral Gyrus | R | 1263 | 51 | 8 | 47 | 29.76 | < 0.001 |
| Triangular Inferior Frontal Gyrus | R |  | 51 | 32 | 5 | 17.25 |  |
| Insula | R |  | 39 | 26 | -1 | 10.18 |  |
| Superior Temporal Gyrus | R | 5507 | 48 | -37 | 5 | 29.41 | < 0.001 |
| Middle Temporal Gyrus | R |  | 51 | -55 | 2 | 28.17 |  |
| Middle Temporal Gyrus | L |  | -48 | -58 | 5 | 25.99 |  |
| Precentral Gyrus | L | 1081 | -48 | 5 | 53 | 20.37 | < 0.001 |
| Precentral Gyrus | L |  | -42 | -1 | 38 | 15.43 |  |
| Orbital Inferior Frontal Gyrus | L |  | -39 | 26 | -4 | 12.00 |  |
| Supplementary Motor Area | R | 674 | 6 | 14 | 56 | 19.62 | < 0.001 |
| Cerebellum 8 | L | 150 | -27 | -64 | -52 | 13.95 | < 0.001 |
| Cerebellum 7b | L |  | -12 | -76 | -43 | 11.97 |  |
| Hippocampus/Thalamus | L | 313 | 18 | -25 | -7 | 13.67 | < 0.001 |
| Hippocampus/Thalamus | R |  | -21 | -25 | -7 | 12.57 |  |
| Vermis 3 | R |  | 3 | -34 | -4 | 9.41 |  |
| Amygdala/Hippocampus | R | 35 | 21 | -7 | -16 | 9.14 | < 0.001 |
| Inferior Parietal Lobule | L | 106 | -24 | -55 | 47 | 8.85 | < 0.001 |
| Amygdala/Hippocampus | L | 23 | -18 | -7 | -16 | 7.47 | < 0.001 |
| Postcentral Gyrus | R | 87 | 48 | -25 | 41 | 6.69 | < 0.001 |
| Postcentral Gyrus | R |  | 57 | -16 | 35 | 5.96 |  |
| Cerebellum Crus2 | R | 14 | 12 | -76 | -40 | 6.41 | < 0.001 |

Notes: All with $p < 0.05$ peak-level FWE correction. Coordinates of peak voxels (x/y/z) are given in Montreal Neurological Institute (MNI) space. L: left, R: right.

**Table S3.** Results from the moderation analyses predicting memory accuracy of fearful faces

| Model | Predictors | B | S.E. | *t* | *p* | R<sup>2</sup> | *F* | *p* |
| --- | --- | --- | --- | --- | --- | --- | --- | --- |
| 1 | Group | -0.0374 | 0.0232 | -1.6066 | 0.1103 | 0.0640 | 1.4465 | 0.1910 |
|  | LAMG | -0.0313 | 0.0280 | -1.1178 | 0.2655 |  |  |  |
|  | Interaction | -0.0249 | 0.0438 | -0.5687 | 0.5704 |  |  |  |
| 2 | Group | -0.0502 | 0.0253 | -1.9853 | 0.0490 | 0.0573 | 1.2856 | 0.2612 |
|  | LAMG-RSTS | -0.0300 | 0.0197 | -1.5259 | 0.1292 |  |  |  |
|  | Interaction | 0.0389 | 0.0244 | 1.5955 | 0.1127 |  |  |  |
| 3 | Group | -0.0709 | 0.0256 | -2.7701 | 0.0063 | 0.0914 | 2.1277 | 0.0441 |
|  | LAMG-LIFG | -0.0453 | 0.0176 | -2.5751 | 0.0110 |  |  |  |
|  | Interaction | 0.0644 | 0.0228 | 2.8284 | 0.0053 |  |  |  |

Notes: All models were adjusted by sex, age, levels of depression and social anxiety. LAMG: activity parameter estimates in the cluster of left amygdala (-24, -7, -22) under imitation. LAMG-RSTS: connectivity parameter estimates in the cluster of right superior temporal sulcus (57, -31, 11) under imitating fearful faces (left amygdala as seed region); LAMG-LIFG: connectivity parameter estimates in the cluster of left inferior frontal gyrus (-42, 26, 2) under imitating fearful faces (left amygdala as seed region).

**Table S4.** Results from the moderation analyses predicting the composite social network scores

| Model | Predictors | B | S.E. | *t* | *p* | R<sup>2</sup> | *F* | *p* |
| --- | --- | --- | --- | --- | --- | --- | --- | --- |
| 1 | Group | -2.9184 | 2.2989 | -1.2694 | 0.2063 | 0.2076 | 5.4274 | <0.001 |
|  | LSTSroi | -2.8300 | 2.3890 | -1.1846 | 0.2381 |  |  |  |
|  | Interaction | 1.6745 | 2.7912 | 0.5999 | 0.5495 |  |  |  |
| 2 | Group | -7.2489 | 2.8700 | -2.5257 | 0.0126 | 0.2270 | 6.0822 | <0.001 |
|  | LAMG-LSTSroi | -1.8602 | 1.8792 | -0.9899 | 0.3239 |  |  |  |
|  | Interaction | 4.8193 | 2.3256 | 2.0723 | 0.0400 |  |  |  |

Notes: All models were adjusted by sex, age, levels of depression and social anxiety. LSTSroi: activity parameter estimates in the predefined ROI of left superior temporal sulcus (-58, -44, 22) across all condition. LAMG-LSTSroi: connectivity parameter estimates in the predefined ROI of left superior temporal sulcus (-58, -44, 22) under imitation (left amygdala (-24, -7, -22) as seed region).

